# Genome-wide association analysis reveals new loci for leaf water status, biomass and plant architectural traits in bread wheat under rainfed conditions

**DOI:** 10.64898/2026.07.11.737917

**Authors:** Samira Rustamova, Atabay Jahangirov, Ulduza Gurbanova, Faig Khudayev, Jens Léon, Ali Ahmad Naz, Irada Huseynova

## Abstract

Drought during reproductive development and grain filling is a major constraint to bread wheat productivity in rainfed environments. In the present study, we employed genome-wide association analysis in an untapped diversity panel of wheat genotypes relevant to natural dryland conditions. A total of 186 genotypes were evaluated for drought-related physiological, biomass, and architectural traits under terminal rainfed stress in Azerbaijan. Relative water content, plant height, fresh weight, dry weight, flag leaf length, and flag leaf width were assessed at the milk ripening stage. These data were subjected to genome-wide association analysis using 19,737 SNP markers to identify loci and epistatic interactions involved in the determination of these traits. The panel showed broad phenotypic variation and significant genotypic effects for all traits, with broad-sense heritability ranging from 0.991 for plant height to 0.385 for flag leaf width. GWAS identified a major locus for relative water content on chromosome 2D at SNP marker AX-86184518, which explained 11.94% of the genotypic variation. Candidate-gene analysis highlighted the proximal WEB-family-like gene TraesCS2D03G1001000 as the main candidate gene. Plant height showed strong additive loci, mainly on chromosomes 2A and 4A, whereas biomass and flag leaf traits showed suggestive additive loci and epistatic interactions. These findings provide candidate loci and interaction patterns in the genetic make-up of essential traits, which may facilitate indirect selection in breeding new varieties.

## 1. Introduction

Bread wheat (*Triticum aestivum* L.) is central to global food security because it contributes substantially to dietary energy, plant-based protein and other nutritional components in human diets (Poole et al., 2021). Recent FAOSTAT data indicate that global wheat production remains close to 800 million tonnes annually from more than 200 million hectares, confirming wheat as one of the most widely grown food crops worldwide (FAO, 2026). Maintaining this production level under changing climatic conditions is becoming increasingly difficult. Heat, drought and compound dry–hot events are expected to occur more frequently during wheat-growing seasons, with particularly strong consequences for rainfed production systems (IPCC, 2023; He et al., 2024). Because a large share of global wheat is produced under rainfed conditions, yield stability is strongly influenced by the timing, duration and intensity of water deficit (Dadrasi et al., 2023). Drought occurring during reproductive development and grain filling is especially damaging, as it restricts canopy function, accelerates senescence and reduces assimilate availability for grain formation (Farooq et al., 2014; Ahmed et al., 2022).

Drought reduces wheat performance through a sequence of linked physiological and growth responses, beginning with impaired plant water status and progressing to reduced carbon assimilation, canopy development and biomass accumulation. Leaf relative water content (RWC) provides a direct estimate of leaf hydration because it relates fresh tissue mass to fully turgid and oven-dry mass (Turner, 1981). In bread wheat, recent morpho-physiological screening under well-watered and drought-stressed conditions confirmed that RWC varies significantly among genotypes and declines under water deficit, while its association with excised leaf water retention, relative water loss and membrane stability reflects broader differences in leaf water status and drought response (Sewore et al., 2023). Water-relation analyses in contrasting wheat genotypes further indicate that drought responsiveness depends on the coordination between leaf water status, stomatal regulation, osmotic adjustment and cell wall elasticity rather than on leaf water content alone (Guizani et al., 2023). Thus, RWC, fresh weight (FW) and dry weight (DW) provide complementary information on the physiological and growth components of drought response, making them suitable traits for genetic analysis under field moisture limitation.

Plant architecture is another important component of dryland adaptation because it influences lodging resistance, biomass distribution, harvest index and suitability to target production environments. Plant height (PH) is not simply a trait to be reduced: excessively tall plants are more prone to lodging, whereas excessively short plants may show reduced biomass and yield potential (Zhang et al., 2025). Recent genetic analysis of plant-height-related traits in wheat also showed that dryland cultivars were approximately 10 cm taller than irrigated cultivars, supporting the view that optimal plant architecture may differ between water-limited and irrigated environments (Zhang et al., 2025). The Green Revolution semi-dwarfing alleles *Rht-B1b* and *Rht-D1b* improved lodging resistance and harvest index, but gibberellin-insensitive reduced-height alleles can shorten coleoptiles and reduce early seedling vigor, which may be disadvantageous under dry sowing or deep-sowing conditions (Rebetzke and Richards, 2000; Hao et al., 2023). Alternative reduced-height loci therefore remain relevant for dryland breeding, particularly when moderate height reduction is needed without compromising early establishment; *Rht25*, for example, has been characterized as a gibberellin-sensitive dwarfing locus affecting PH in wheat (Mo et al., 2018). In addition to plant height, flag leaf length (FLL) and flag leaf width (FLW) are agronomically relevant because the flag leaf is a major photosynthetic source during grain filling. Genetic dissection of flag leaf morphology under contrasting water regimes has shown that these traits are quantitatively inherited and responsive to water availability (Yang et al., 2016). Flag-leaf-related traits, including length and width, have also been associated with plant architecture and grain yield in wheat, supporting their value as target traits in genetic analysis (Wang et al., 2022). More recently, natural allelic variation has been shown to contribute to diversity in the regulation of wheat flag leaf morphology, reinforcing the value of these traits for association mapping in diverse germplasm (Schierenbeck et al., 2024). However, the extent to which flag leaf morphology is genetically coordinated with foliar water status and biomass accumulation under rainfed field stress remains insufficiently resolved.

The genetic dissection of drought-related traits is complicated by their quantitative inheritance and sensitivity to environmental conditions. Drought adaptation in wheat is generally controlled by multiple physiological, developmental and genetic mechanisms rather than by single major-effect loci (Sallam et al., 2019). Biparental quantitative trait locus (QTL) mapping has been useful for detecting segregating loci in specific populations, but its allelic coverage is limited by the two parental genotypes used in each cross. Genome-wide association stady (GWAS) provides a complementary approach because it exploits historical recombination and linkage disequilibrium in diverse panels to detect marker–trait associations at higher resolution. In wheat, GWAS has been used to identify loci associated with seedling drought tolerance (Maulana et al., 2020), osmotic adjustment and related drought-adaptive traits in durum wheat (Condorelli et al., 2022), and drought-induced proline and hydrogen peroxide accumulation under natural field and rain-out shelter conditions (Kamruzzaman et al., 2022). In Turkish winter wheat germplasm, the integration of GWAS and selective sweep analysis identified genomic regions and candidate genes associated with improved grain yield under drought conditions (Sehgal et al., 2024). The biological interpretation of these associations has been strengthened by the availability of annotated wheat genome resources, including the reference genome and multi-genome assemblies (IWGSC, 2018; Walkowiak et al., 2020). More recent studies have identified single-nucleotide polymorphism (SNP) markers, candidate genes and gene networks associated with drought tolerance and drought-response indices in wheat (Nouraei et al., 2024; Mosalam et al., 2025; Mourad et al., 2026). However, most GWAS studies still emphasize additive marker effects, whereas non-additive marker interactions may also contribute to trait variation under field stress. Therefore, combining additive GWAS with epistatic interaction analysis may provide a more complete view of the genetic basis of drought-related variation in adapted wheat germplasm.

Despite increasing progress in wheat drought genetics, important knowledge gaps remain for rainfed field environments. Many studies have used seedling-stage assays, hydroponic or PEG-induced stress, or controlled drought treatments that do not fully reproduce the fluctuating moisture availability, temperature conditions and soil constraints experienced in dryland production systems. In addition, the genetic coordination among foliar hydration, biomass accumulation and plant architecture remains insufficiently understood in locally and regionally adapted germplasm evaluated under real field moisture limitation. Addressing this gap is important because drought adaptation in breeding programs depends not only on statistically significant markers, but also on markers associated with biologically interpretable traits expressed in target production environments.

Being integral geographic part of wheat center of diversity and origin, the dryland conditions of Azerbaijan provide a relevant field context for rainfed wheat production environment globally. Under these natural dryland conditions, the production is exposed to variable precipitation and terminal-season moisture limitation, while local and regionally adapted germplasm may retain allelic variation shaped by recurrent water-limited conditions (Red Cross Red Crescent Climate Centre, 2024). Previous studies on Azerbaijani wheat germplasm have revealed substantial variation in drought-responsive physiological, biochemical, ultrastructural and molecular traits (Rustamova et al. 2021; Aliyeva et al., 2023; Allahverdiyev, 2024; Isgandarova et al., 2024; Allahverdiyev et al. 2025). However, the genome-wide basis of this adaptive variation remains insufficiently characterized under rainfed field conditions.

The present study evaluated a diverse panel of 186 bread wheat genotypes under rainfed field conditions in Azerbaijan. The objectives were to: (1) characterize phenotypic variation, broad-sense heritability and inter-trait relationships for RWC, PH, FW, DW, FLL and FLW under rainfed field conditions; (2) identify additive marker–trait associations using high-density SNP-based GWAS; (3) evaluate epistatic marker–marker interactions contributing to drought-related trait variation; and (4) identify candidate genes located near significant marker. By linking physiological, architectural and biomass traits with genome-wide marker information, this study aims to clarify the genetic basis of drought-related variation in field-grown wheat and to support the identification of markers and candidate genes relevant for breeding in water-limited environments.

## 2. Materials and Methods

### 2.1. Plant material

The plant material utilized in the present study consists of a newly established diversity set of 186 genotypes of bread wheat (Triticum aestivum L.) adapted to natural rainfed conditions representing a wide geographic regions across the center of wheat diversity and its origin (Supplementary Table 1). This collection was designed to capture a broad range of phenotypic and genetic diversity relevant for drought adaptation under rainfed conditions.

### 2.2. Experimental design and growth condition

Field experiments were conducted during the 2022/2023 growing season at the Gobustan Regional Experimental Station of the Research Institute of Crop Husbandry, Azerbaijan (800 m a.s.l.). The soil in this region is classified as light chestnut with a slightly alkaline pH. It is characterized by low humus content (1.5-1.8% in the topsoil), low nitrogen and phosphorus availability, and moderate potassium levels. The hydro-meteorological conditions for the 2022/2023 growing season are presented in detail in Supplementary Table 2. Total precipitation during the cropping season was 356.3 mm, which is below the long-term average of 406.0 mm. A critical moisture deficit occurred in June, with only 4.1 mm of rainfall accompanied by relatively high temperatures, causing pronounced drought stress during the reproductive stages of wheat development. Each experimental plot measured 1.5 × 1.0 m (1.5 m²), with rows spaced 15 cm apart. Sowing was performed at a density of 450 germinating seeds per m². The experiment followed a randomized complete block design with three replications per genotype.

### 2.3. Phenotypic evaluation of drought-related traits

Phenotypic measurements were conducted during the first ten days of June 2023, at the milk ripeness stage. Six drought-related agronomic and physiological traits were evaluated: PH, FLL, FLW, FW, DW, and RWC.

PH was measured from the soil surface to the top of the spike, excluding awns. FLL was measured from the base to the tip of the fully developed flag leaf, while FLW was measured at the widest part of the same leaf using a ruler (Yang et al., 2016).

For fresh and dry weight determination, leaf samples were collected from each genotype and replication. FW was recorded immediately after sampling. The samples were then dried in a thermostat at 80°C until constant weight, and DW was recorded.

RWC was measured according to the method proposed by Turner (1981). Fresh, medium-sized leaves were weighed immediately after collection to determine FW. The leaves were then immersed in distilled water for 24 h to reach full turgidity, after which turgid weight (TW) was recorded. Subsequently, the samples were dried at 80°C until constant weight and weighed again to determine DW. RWC was calculated using the following formula:

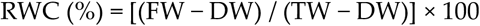

where FW is the initial fresh weight of the leaf, TW is the turgid weight after rehydration, and DW is the dry weight after oven-drying.

### 2.4. Broad sense heritability (H2)

Descriptive statistics were calculated using R software (v3.6). A mixed linear model (MLM) and two-way ANOVA were performed using the ’nlme’ and ’emmeans’ packages to estimate variance components and compare means, considering cultivar and replication as random effects. The variance components were used for calculating broad sense heritability (H2). The following formula was used for estimating H2:

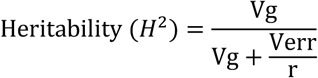

Where, Vg is genotypic variance and Verr is the error variance and r is the number of replications.

### 2.5. Pearson correlation analysis

Pearson correlation analysis was performed to evaluate the relationships among the six drought-related traits, including RWC, PH, DW, FW, FLL, and FLW. Correlation coefficients were calculated using trait mean values across genotypes. The resulting correlation matrix was visualized as a heatmap to identify coordinated variation among physiological, biomass-related, and plant-architecture traits under rainfed field conditions.

### 2.6. Genotyping of the diversity-set

The wheat accessions were genotyped by TraitGenetics GmbH (Gatersleben, Germany) using the Illumina Infinium XT 25K SNP array. An initial set of 24,146 SNPs was subjected to quality control, which included the removal of monomorphic markers, SNPs with missing data, and markers with a minor allele frequency (MAF) below 5%. This filtering resulted in a final dataset of 19,737 high-quality polymorphic SNPs used for all downstream analyses. Details of these markers are described in Zakieh et al. (2021, 2023).

### 2.7. GWAS

Genome-wide association mapping was performed following the GRAMMAR method described by Aulchenko et al. (2007). For the analysis, the first three principal components were included as co-factors to control for population structure in the analysis additionally. The methods of the analysis were used and described in detail by Reinert et al. (2016) and Naz et al. (2017). We used a linear mixed model to calculate the QTLs as presented below:

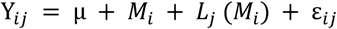

where Yij is the phenotypic value; μ is the general mean; Mi is the fixed effect of ith marker genotype; Lj (Mi) is the random effect of j-th Cultivar nested within i-th marker genotype and εij is the residual. Significant markers were detected following a correction using the probability of false discovery rate (FDR), implemented in the SAS procedure PROC MULTTEST according to Benjamini and Yekutieli (2005). To account for multiple testing across all SNPs, genome-wide significance was additionally assessed using a Bonferroni correction based on the total number of markers tested (P ≤ 2.53 × 10⁻⁶).

### 2.8. Epistatic interaction analysis

Pairwise epistatic marker interactions were analyzed using a two-way multilocus approach implemented in the PROC MIXED procedure of SAS, following the general modelling strategy described by Benaouda et al. (2022). To reduce the computational burden of testing all possible marker-by-marker combinations, only markers showing an initial single-marker association signal of P < 0.7 were retained for epistasis testing.

Because the present study was conducted in a single field environment, the environmental interaction term was not included in the model. The following model was used:

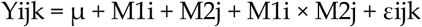

where Yijk is the phenotypic value, μ is the general mean, M1i is the fixed effect of the i-th genotype class of the first marker, M2j is the fixed effect of the j-th genotype class of the second marker, M1i × M2j is the fixed interaction effect between the two marker genotype classes, and εijk is the residual error.

Epistatic interactions were evaluated for all six drought-related traits: RWC, PH, DW, FW, FLL, and FLW. Marker-by-marker interaction effects were assessed based on the probability value of the interaction term. For each selected marker pair, allele-combination means were calculated to describe the phenotypic effect associated with different two-locus genotype combinations.

Reciprocal duplicate pairs, in which the same marker combination appeared as Marker A × Marker B and Marker B × Marker A, were collapsed into a single unique marker pair. For each retained interaction, the minimum and maximum allele-combination means were determined, and the mean range was calculated as the difference between the highest and lowest allele-combination mean. The strongest representative interactions were ranked by probability value and summarized in the main text, while the complete list of epistatic marker combinations was provided in the supplementary materials. Because the analysis was performed in a single field environment, these interactions were interpreted as putative non-additive signals requiring further validation.

### 2.9. Determination of genes colocalizing with QTL

To identify putative candidate genes underlying significant QTL, lead SNP markers were anchored to the wheat reference genome (IWGSC RefSeq v2.1) based on their physical positions in the Ensembl-plant database (Bolser et al. 2017). Candidate gene mining was performed using the BioMart interface (Smedley et al. 2015). A fixed-window approach was applied by defining a ±500 kb genomic interval on both sides of each lead SNP. All annotated gene models whose start or end coordinates fell within this interval were retrieved and considered as putative candidate genes. Functional annotation of candidate genes was obtained based on Gene Ontology (GO) terms describing molecular functions and biological processes, using information available in the UniProt database (The UniProt Consortium 2023).

### 2.10. In silico expression analysis of candidate genes

To support the functional interpretation of genes located within candidate QTL intervals, publicly available wheat transcriptome data were inspected using the Wheat Expression Browser, which is based on the expVIP platform (Borrill et al., 2016) and the wheat expression atlas described by Ramírez-González et al. (2018). For candidate genes identified in IWGSC RefSeq v2.1, corresponding RefSeq v1.1 gene models were used when available, because the expression browser is primarily organized according to Ref-Seq v1.1 annotations. Expression profiles were examined across available non-stress and abiotic-stress-related samples. Transcript abundance was expressed as transcripts per million (TPM), and mean TPM values were compared descriptively between abiotic-stress-associated and non-abiotic sample groups. This analysis was used only as supportive evidence for prioritizing positional candidate genes and was not treated as experimental validation of gene function.

## 3. Results

### 3.1. Phenotypic variation, heritability and inter-trait correlations

Substantial phenotypic variation was observed among the 186 bread wheat genotypes for all six drought-related traits under rainfed field conditions (Table 1). RWC ranged from 41.14% to 95.97%, with a mean value of 68.15%, indicating broad variation in plant water status under terminal drought. PH varied from 65.00 to 135.33 cm, whereas DW and FW ranged from 0.05 to 0.29 g and from 0.17 to 1.03 g, respectively. FLL ranged from 14.00 to 31.93 cm, while FLW varied from 1.00 to 2.70 cm.

**Table 1.** Phenotypic variation and broad-sense heritability of drought-related traits under rainfed field conditions.

| Traits | Max | Min | Mean | CV (%) | H<sup>2</sup> | Genotype |
| --- | --- | --- | --- | --- | --- | --- |
| RWC | 95.97 | 41.14 | 68.15 | 13.74 | 0.639 | *** |
| PH | 135.33 | 65.00 | 97.70 | 13.80 | 0.991 | *** |
| DW | 0.29 | 0.05 | 0.12 | 31.39 | 0.785 | *** |
| FW | 1.03 | 0.17 | 0.47 | 31.02 | 0.775 | *** |
| FLL | 31.93 | 14.00 | 21.10 | 14.53 | 0.654 | *** |
| FLW | 2.70 | 1.00 | 1.76 | 14.12 | 0.385 | *** |

\*H<sup>2</sup>, broad-sense heritability; CV, coefficient of variation; RWC, relative water content; PH, plant height; DW, dry weight; FW, fresh weight; FLL, flag leaf length; FLW, flag leaf width. RWC is expressed as percentage; PH, FLL, and FLW are expressed in cm; FW and DW are expressed in g. The genotype column indicates the significance of genotypic effects based on analysis of variance. \*\*\*, $P < 0.001$ .

The coefficient of variation was highest for the two biomass-related traits, DW (31.39%) and FW (31.02%), suggesting greater relative dispersion for biomass accumulation under rainfed conditions. In contrast, RWC, PH, FLL and FLW showed moderate coefficients of variation, ranging from 13.74% to 14.53%. Genotypic effects were highly significant for all traits (P < 0.001), confirming the presence of substantial genetic variation within the panel.

Broad-sense heritability differed considerably among traits. The highest heritability was observed for PH (H² = 0.991), followed by DW (H² = 0.785), FW (H² = 0.775), FLL (H² = 0.654), RWC (H² = 0.639), and FLW (H² = 0.385). These results indicate that PH and bio-mass-related traits were under stronger genetic control in the present field environment, whereas FLW was more strongly influenced by environmental variation or genotype-by-environment sensitivity.

Pearson’s correlation analysis was performed to evaluate the relationships among physiological, biomass-related, and architectural traits (Figure 1). The strongest positive correlation was observed between FW and DW (*r* = 0.88, *P* < 0.001), indicating a close relationship between fresh and dry biomass accumulation. FW was also strongly and positively correlated with FLL (*r* = 0.75, *P* < 0.001) and FLW (*r* = 0.70, *P* < 0.001), while DW showed positive correlations with FLW (*r* = 0.69, *P* < 0.001) and FLL (*r* = 0.64, *P* < 0.001). A moderate positive correlation was detected between FLL and FLW (*r* = 0.35, *P* < 0.001), suggesting partial coordination between the two flag leaf architectural traits.

**Figure 1.**
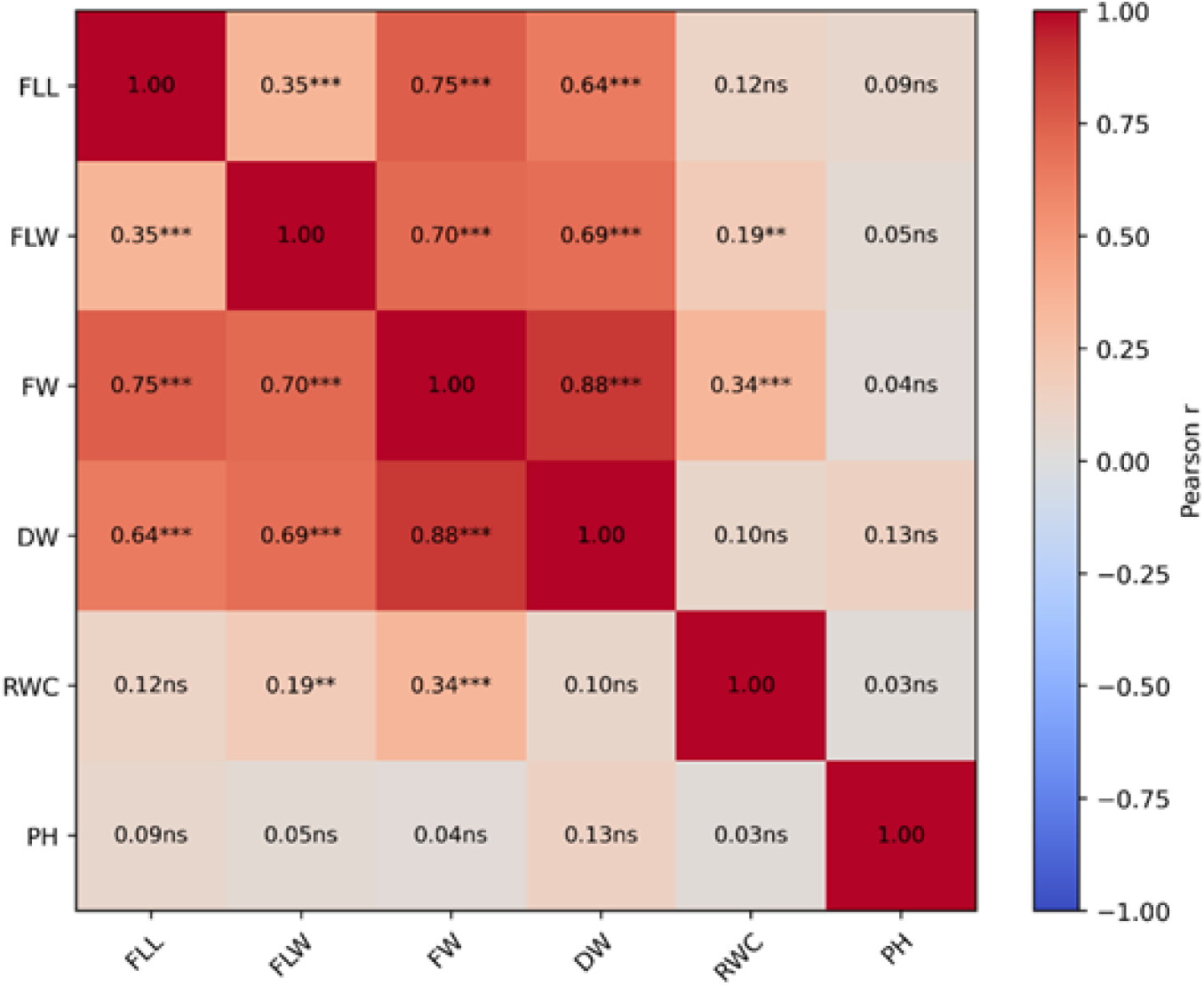
Pearson correlation matrix of drought-related traits in bread wheat under rainfed field conditions. Pearson correlation coefficients (r) were calculated among flag leaf length (FLL), flag leaf width (FLW), fresh weight (FW), dry weight (DW), relative water content (RWC), and plant height (PH). The heatmap shows the strength of pairwise phenotypic correlations among traits; color intensity reflects the magnitude of the correlation coefficient. Asterisks indicate statistical significance: P < 0.01 (**), and P < 0.001 (***), while “ns” indicates non-significant correlations.

RWC showed a moderate positive correlation with FW (*r* = 0.34, *P* < 0.001) and a weaker but significant positive correlation with FLW (*r* = 0.19, *P* < 0.01). In contrast, RWC was not significantly correlated with FLL (*r* = 0.12), DW (*r* = 0.10), or PH (*r* = 0.03). PH also showed no significant correlations with the other measured traits, including FLL (*r* = 0.09), FLW (*r* = 0.05), FW (*r* = 0.04), DW (*r* = 0.13), and RWC (*r* = 0.03). Overall, the correlation structure suggests that biomass accumulation and flag leaf morphology were closely associated, whereas PH behaved largely independently from the other drought-related traits in this panel. The partial association of RWC with FW and FLW further indicates that foliar water status was related to some, but not all, components of biomass accumulation and leaf architecture under rainfed conditions.

### 3.2. Marker–trait associations using GWAS and multiple-testing correction

Genome-wide association analysis was performed using 19,737 high-quality polymorphic SNP markers. The strength and number of marker–trait associations differed markedly among the six evaluated traits (Supplementary Table 3). At the suggestive threshold of *P* < 1 × 10⁻⁴, PH showed the largest number of associated SNPs, with 114 SNPs detected. RWC showed ten SNPs with *P* < 1 × 10⁻⁴, followed by FLW with nine SNPs, DW with four SNPs, FW with two SNPs, and FLL with one SNP.

After multiple-testing correction, the strongest statistical evidence was observed for PH and RWC. PH included SNPs passing both FDR and Bonferroni thresholds, whereas RWC included one SNP that passed both corrected thresholds. In contrast, DW, FW, FLL, and FLW showed suggestive SNP-level associations but no associations passing the applied FDR or Bonferroni thresholds (Supplementary Table 3). Therefore, associations detected for these traits were interpreted as preliminary statistically identified marker–trait effects.

The Manhattan plots illustrated this trait-specific pattern, with the most pronounced associations observed for PH, one corrected association for RWC, and mainly suggestive marker–trait associations for biomass and flag leaf traits (Figure 2). In these plots, red points indicate peak markers retained after marker selection based on association strength (*P* < 1 × 10⁻⁴) and local linkage disequilibrium, whereas blue points represent the remaining tested SNPs. Based on these results, subsequent interpretation was structured according to both biological trait groups and statistical evidence level, distinguishing corrected SNP associations from suggestive SNP-level associations.

**Figure 2.**
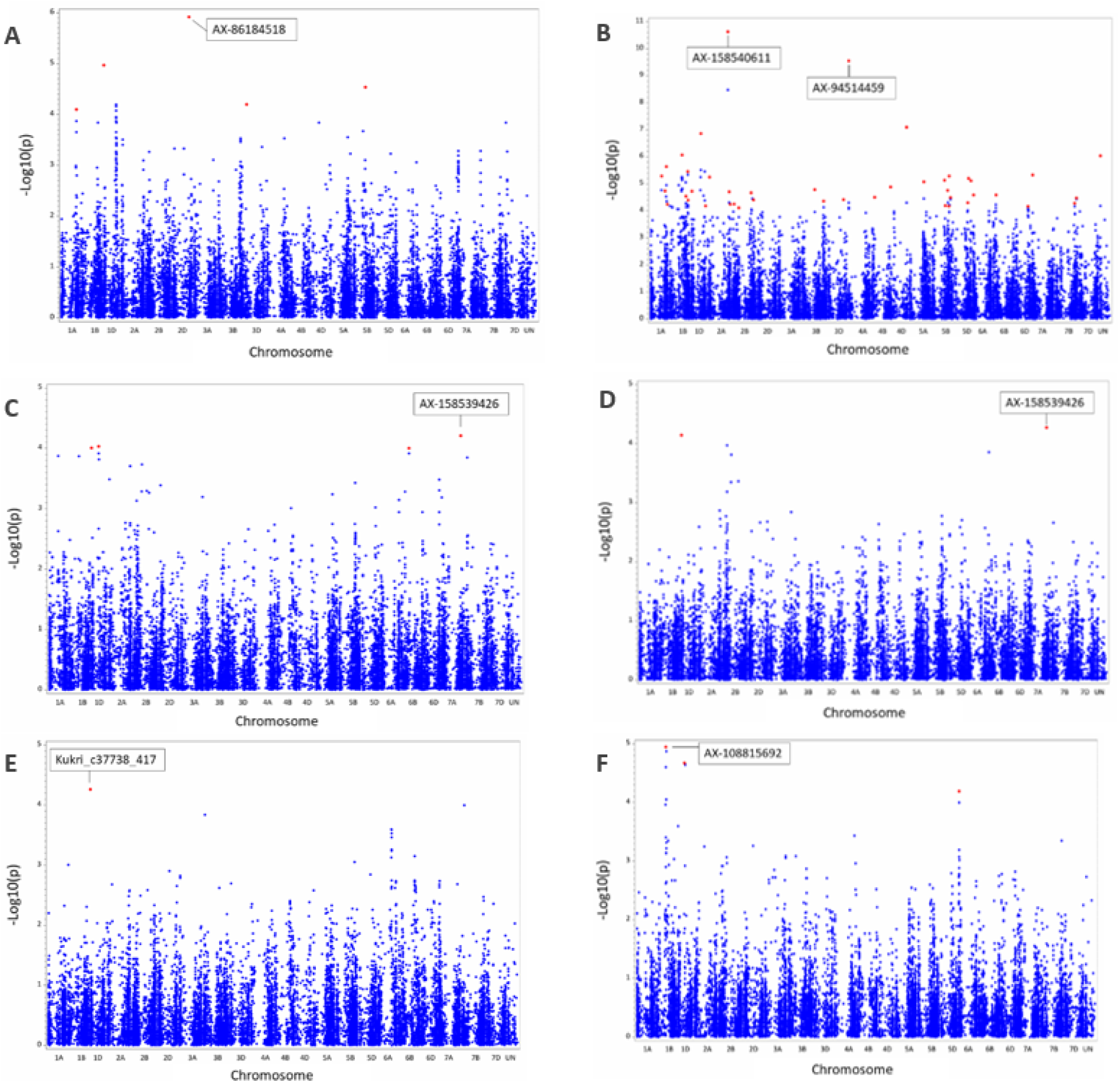
Manhattan plots of genome-wide SNP marker–trait associations for drought-related traits under rainfed field conditions. Manhattan plots are shown for relative water content (RWC; A), plant height (PH; B), dry weight (DW; C), fresh weight (FW; D), flag leaf length (FLL; E), and flag leaf width (FLW; F). Each point represents a SNP plotted according to its chromosomal position and −log₁₀(P) value. Red points indicate peak markers retained after marker selection based on association strength (P < 1 × 10⁻⁴) and local linkage disequilibrium (LD), whereas blue points represent the remaining tested SNPs.

### 3.3 Marker–trait associations for RWC

RWC was considered a key physiological indicator of plant water status under rainfed conditions. GWAS identified one corrected significant association for RWC (Table 2). The strongest RWC-associated SNP was AX-86184518 on chromosome 2D, located at 560,005,079 bp (Figure 2A). This marker showed P = 1.21 × 10⁻⁶ and explained 11.94% of the genotypic variation. The two allele classes differed substantially in mean RWC values, with 67.75% for the major allele class and 82.00% for the minor allele class, suggesting a marked marker-associated difference in plant water status. Additional suggestive RWC-associated SNPs were detected on chromosomes 1D, 5B and 3D (Table 2). These SNPs did not pass the corrected significance threshold and were therefore interpreted cautiously. Among all RWC signals, the chromosome 2D region tagged by AX-86184518 represented the most robust association detected in the present analysis.

**Table 2.**
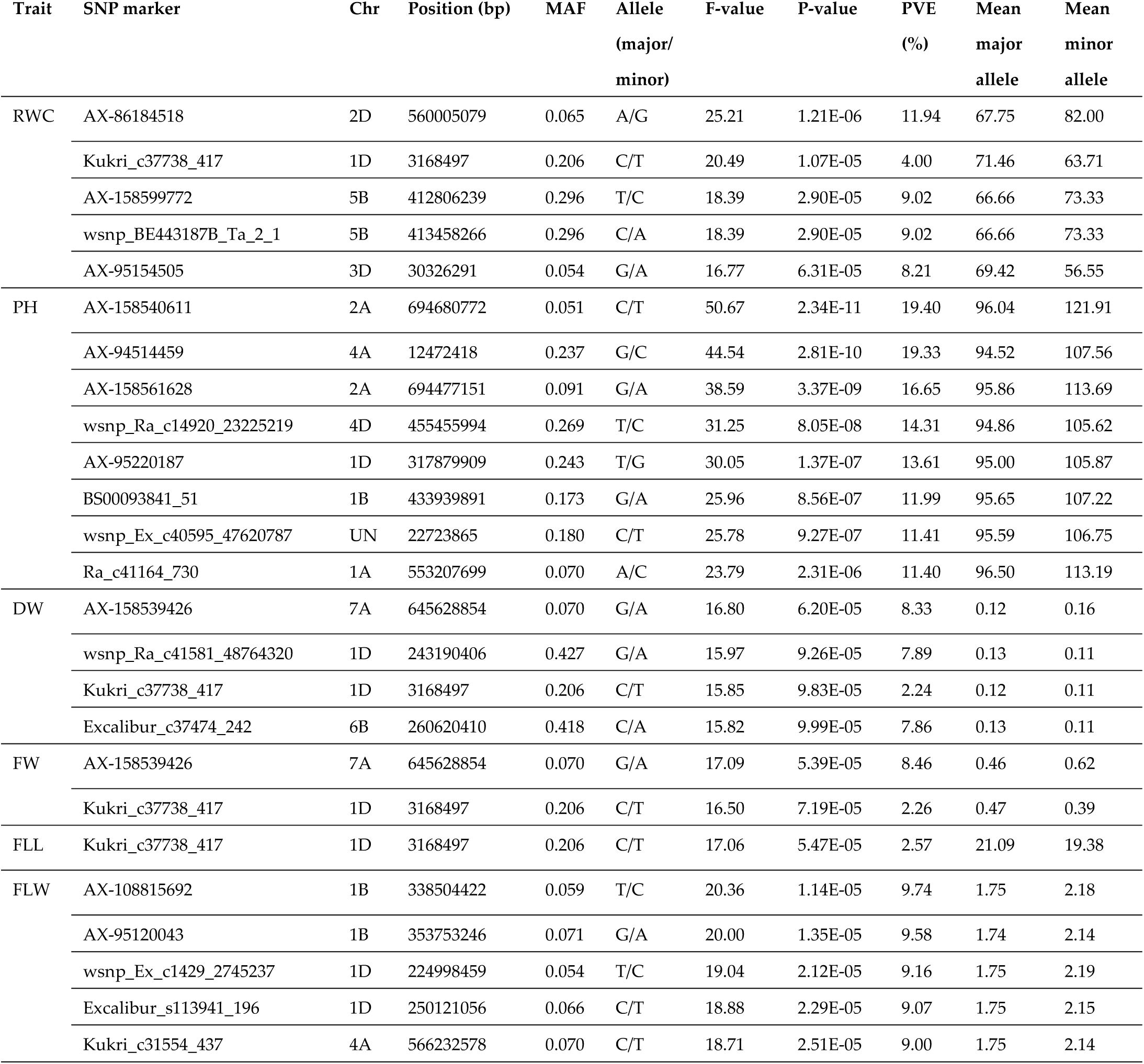
Selected SNP marker–trait associations detected for drought-related traits.

### 3.4. Marker–trait associations for plant growth and biomass-related traits

Among the analyzed growth-related traits, PH showed the strongest evidence for additive marker–trait associations. At the suggestive threshold of *P* < 1 × 10⁻⁴, 114 SNPs were detected for PH (Supplementary Table 3). Multiple-testing correction further supported PH as the trait with the strongest GWAS evidence, with several SNPs passing both FDR and Bonferroni thresholds. The leading PH association was detected on chromosome 2A at marker AX-158540611, located at 694,680,772 bp (Figure 2B). This marker showed *P* = 2.34 × 10⁻¹¹ and explained 19.40% of the genotypic variation (Table 2). The mean PH values differed strongly between allele classes, with 96.04 cm for the major allele class and 121.91 cm for the minor allele class. A second major PH-associated SNP was detected on chromosome 4A at marker AX-94514459, located at 12,472,418 bp (Figure 2B). This marker showed *P* = 2.81 × 10⁻¹⁰ and explained 19.33% of the genotypic variation. Additional Bon-ferroni-significant PH-associated SNPs were located on chromosomes 2A, 4D, 1D, 1B, and 1A, as well as in one unassigned genomic region (Table 2). Together, these results indicate that PH in this wheat panel was associated with multiple genomic regions with relatively strong additive effects.

In contrast, DW and FW showed high broad-sense heritability but did not reveal associations passing the applied multiple-testing thresholds. DW had four SNPs with *P* < 1 × 10⁻⁴. The strongest DW-associated SNP was AX-158539426 on chromosome 7A, located at 645,628,854 bp (Figure 2C). This marker showed *P* = 6.20 × 10⁻⁵ and explained 8.33% of the genotypic variation. Additional suggestive DW-associated SNPs were detected on chromosomes 1D and 6B (Table 2). FW showed two suggestive SNP-level associations, including AX-158539426 on chromosome 7A (Figure 2D) and Kukri_c37738_417 on chromosome 1D. AX-158539426 showed *P* = 5.39 × 10⁻⁵ and explained 8.46% of the genotypic variation, while Kukri_c37738_417 showed *P* = 7.19 × 10⁻⁵ and explained 2.26% of the genotypic variation (Table 2). Because these associations did not pass the corrected significance thresholds, they were interpreted as preliminary statistically identified marker–trait effects.

### 3.5. Marker–trait associations suggested additive associations for flag leaf architecture

Flag leaf architecture was evaluated using FLL and FLW. FLL showed moderate-to-high heritability and one suggestive SNP-level association with *P* < 1 × 10⁻⁴ (Table 2). This association corresponded to marker Kukri_c37738_417 on chromosome 1D, with *P* = 5.47 × 10⁻⁵ and 2.57% of the genotypic variation explained (Figure 2E). Because this marker did not pass the corrected significance threshold, the additive GWAS evidence for FLL was limited to a suggestive marker–trait association.

FLW showed the lowest heritability among the analyzed traits, but nine SNPs reached *P* < 1 × 10⁻⁴ (Supplementary Table 3). The strongest FLW-associated SNP was AX-108815692 on chromosome 1B, located at 338,504,422 bp, with *P* = 1.14 × 10⁻⁵ and 9.74% of the genotypic variation explained (Figure 2F). Additional suggestive FLW-associated SNPs were detected on chromosomes 1B, 1D, and 4A. Among these, AX-95120043 on chromosome 1B explained 9.58% of the genotypic variation, wsnp_Ex_c1429_2745237 on chromosome 1D explained 9.16%, Excalibur_s113941_196 on chromosome 1D explained 9.07%, and Kukri_c31554_437 on chromosome 4A explained 9.00% (Table 2). Although these markers did not pass the corrected significance threshold, they represent preliminary statistically identified marker–trait associations with relatively high explained genotypic variation. Overall, flag leaf architecture traits showed less corrected additive GWAS evidence than PH and RWC, suggesting a more complex or environmentally sensitive genetic architecture.

### 3.6 Putative epistatic marker interactions across drought-related traits

Putative epistatic interaction analysis identified marker-by-marker interactions for all six drought-related traits. After collapsing reciprocal duplicate pairs, the number of unique epistatic marker pairs differed among traits, with 755 pairs for RWC, 754 for DW, 752 for FW, 317 for FLL, 662 for FLW, and 1454 for PH (Supplementary Table 4). These results suggest that the extent of putative non-additive marker interactions varied among the analyzed traits. For phenotypic interpretation, allele-combination means were calculated for the representative marker pairs retained in Table 3. The five top-ranked representative interactions per trait are presented in Table 3, while detailed allele-combination means for these representative interactions are provided in Supplementary Table 5.

**Table 3.**
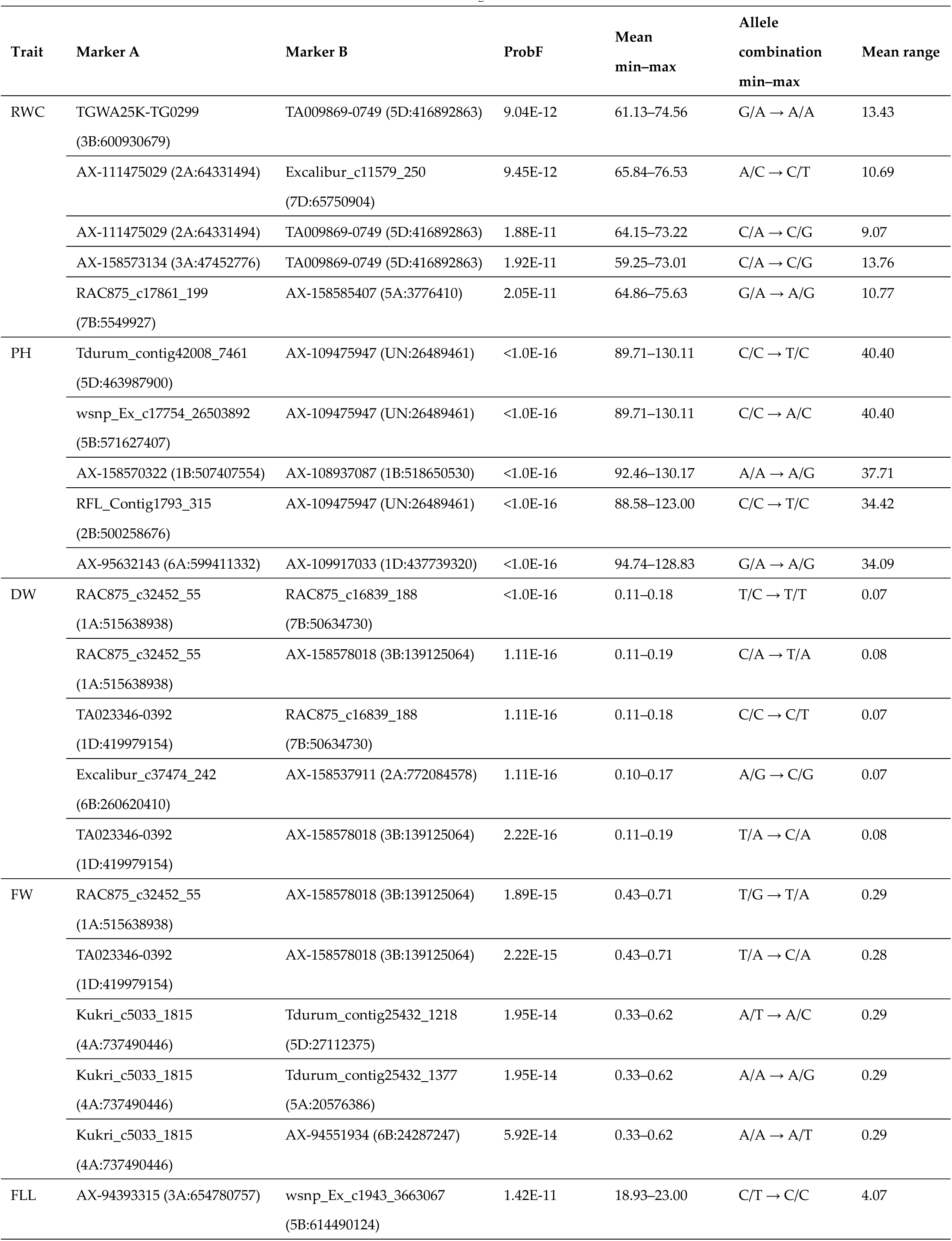
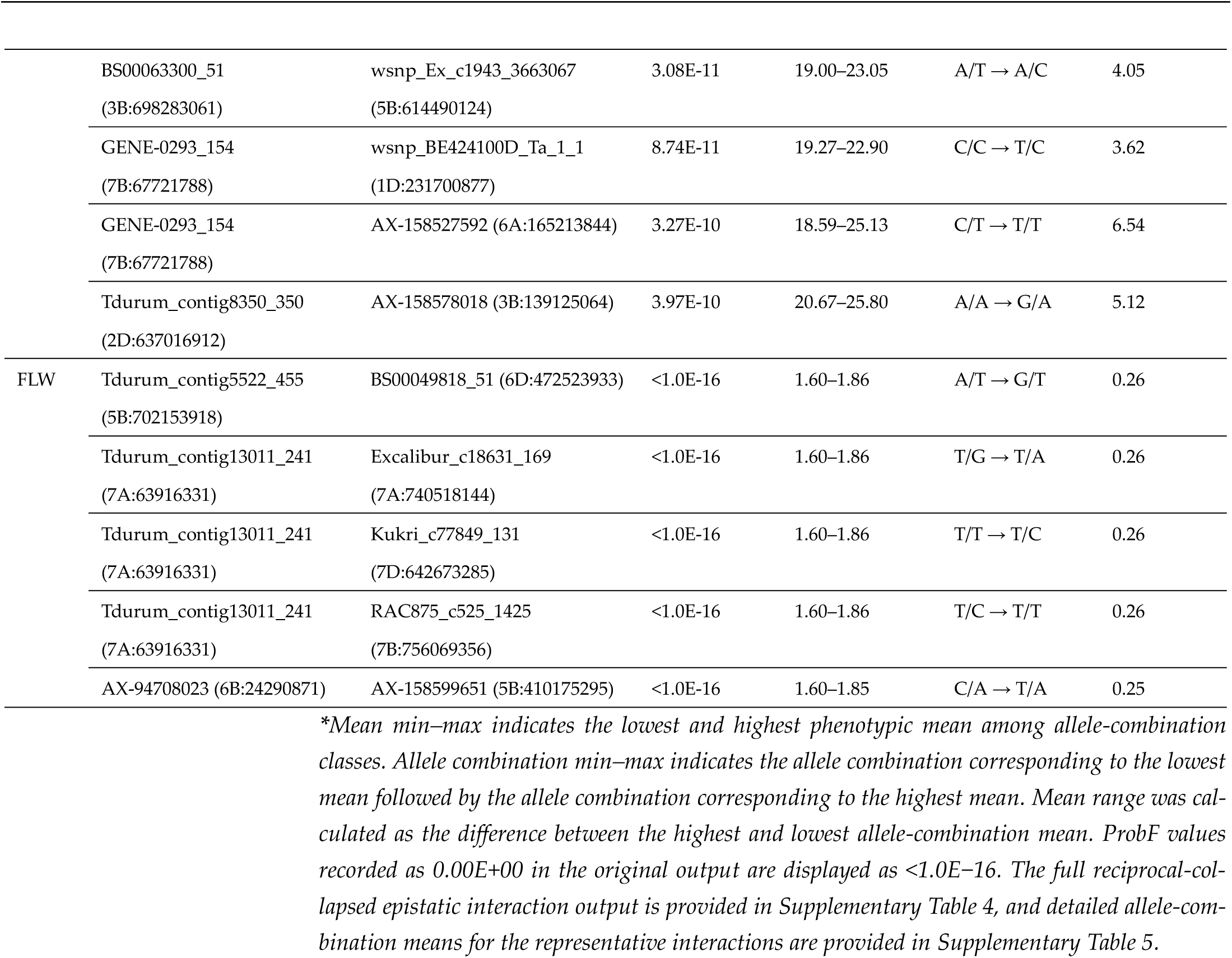
Putative epistatic marker interactions and allele-combination means in the determination of evaluated drought-related traits.

| Trait | Marker A | Marker B | ProbF | Mean<br>min-max | Allele<br>combination<br>min-max | Mean range |
| --- | --- | --- | --- | --- | --- | --- |
| RWC | TGWA25K-TG0299<br>(3B:600930679) | TA009869-0749 (5D:416892863) | 9.04E-12 | 61.13–74.56 | G/A → A/A | 13.43 |
|  | AX-111475029 (2A:64331494) | Excalibur_c11579_250<br>(7D:65750904) | 9.45E-12 | 65.84–76.53 | A/C → C/T | 10.69 |
|  | AX-111475029 (2A:64331494) | TA009869-0749 (5D:416892863) | 1.88E-11 | 64.15–73.22 | C/A → C/G | 9.07 |
|  | AX-158573134 (3A:47452776) | TA009869-0749 (5D:416892863) | 1.92E-11 | 59.25–73.01 | C/A → C/G | 13.76 |
|  | RAC875_c17861_199<br>(7B:5549927) | AX-158585407 (5A:3776410) | 2.05E-11 | 64.86–75.63 | G/A → A/G | 10.77 |
| PH | Tdurum_contig42008_7461<br>(5D:463987900) | AX-109475947 (UN:26489461) | <1.0E-16 | 89.71–130.11 | C/C → T/C | 40.40 |
|  | wsnp_Ex_c17754_26503892<br>(5B:571627407) | AX-109475947 (UN:26489461) | <1.0E-16 | 89.71–130.11 | C/C → A/C | 40.40 |
|  | AX-158570322 (1B:507407554) | AX-108937087 (1B:518650530) | <1.0E-16 | 92.46–130.17 | A/A → A/G | 37.71 |
|  | RFL_Contig1793_315<br>(2B:500258676) | AX-109475947 (UN:26489461) | <1.0E-16 | 88.58–123.00 | C/C → T/C | 34.42 |
|  | AX-95632143 (6A:599411332) | AX-109917033 (1D:437739320) | <1.0E-16 | 94.74–128.83 | G/A → A/G | 34.09 |
| DW | RAC875_c32452_55<br>(1A:515638938) | RAC875_c16839_188<br>(7B:50634730) | <1.0E-16 | 0.11–0.18 | T/C → T/T | 0.07 |
|  | RAC875_c32452_55<br>(1A:515638938) | AX-158578018 (3B:139125064) | 1.11E-16 | 0.11–0.19 | C/A → T/A | 0.08 |
|  | TA023346-0392<br>(1D:419979154) | RAC875_c16839_188<br>(7B:50634730) | 1.11E-16 | 0.11–0.18 | C/C → C/T | 0.07 |
|  | Excalibur_c37474_242<br>(6B:260620410) | AX-158537911 (2A:772084578) | 1.11E-16 | 0.10–0.17 | A/G → C/G | 0.07 |
|  | TA023346-0392<br>(1D:419979154) | AX-158578018 (3B:139125064) | 2.22E-16 | 0.11–0.19 | T/A → C/A | 0.08 |
| FW | RAC875_c32452_55<br>(1A:515638938) | AX-158578018 (3B:139125064) | 1.89E-15 | 0.43–0.71 | T/G → T/A | 0.29 |
|  | TA023346-0392<br>(1D:419979154) | AX-158578018 (3B:139125064) | 2.22E-15 | 0.43–0.71 | T/A → C/A | 0.28 |
|  | Kukri_c5033_1815<br>(4A:737490446) | Tdurum_contig25432_1218<br>(5D:27112375) | 1.95E-14 | 0.33–0.62 | A/T → A/C | 0.29 |
|  | Kukri_c5033_1815<br>(4A:737490446) | Tdurum_contig25432_1377<br>(5A:20576386) | 1.95E-14 | 0.33–0.62 | A/A → A/G | 0.29 |
|  | Kukri_c5033_1815<br>(4A:737490446) | AX-94551934 (6B:24287247) | 5.92E-14 | 0.33–0.62 | A/A → A/T | 0.29 |
| FLL | AX-94393315 (3A:654780757) | wsnp_Ex_c1943_3663067<br>(5B:614490124) | 1.42E-11 | 18.93–23.00 | C/T → C/C | 4.07 |
|  | BS00063300_51<br>(3B:698283061) | wsnp_Ex_c1943_3663067<br>(5B:614490124) | 3.08E-11 | 19.00–23.05 | A/T → A/C | 4.05 |
|  | GENE-0293_154<br>(7B:67721788) | wsnp_BE424100D_Ta_1_1<br>(1D:231700877) | 8.74E-11 | 19.27–22.90 | C/C → T/C | 3.62 |
|  | GENE-0293_154<br>(7B:67721788) | AX-158527592 (6A:165213844) | 3.27E-10 | 18.59–25.13 | C/T → T/T | 6.54 |
|  | Tdurum_contig8350_350<br>(2D:637016912) | AX-158578018 (3B:139125064) | 3.97E-10 | 20.67–25.80 | A/A → G/A | 5.12 |
| FLW | Tdurum_contig5522_455<br>(5B:702153918) | BS00049818_51 (6D:472523933) | <1.0E-16 | 1.60–1.86 | A/T → G/T | 0.26 |
|  | Tdurum_contig13011_241<br>(7A:63916331) | Excalibur_c18631_169<br>(7A:740518144) | <1.0E-16 | 1.60–1.86 | T/G → T/A | 0.26 |
|  | Tdurum_contig13011_241<br>(7A:63916331) | Kukri_c77849_131<br>(7D:642673285) | <1.0E-16 | 1.60–1.86 | T/T → T/C | 0.26 |
|  | Tdurum_contig13011_241<br>(7A:63916331) | RAC875_c525_1425<br>(7B:756069356) | <1.0E-16 | 1.60–1.86 | T/C → T/T | 0.26 |
|  | AX-94708023 (6B:24290871) | AX-158599651 (5B:410175295) | <1.0E-16 | 1.60–1.85 | C/A → T/A | 0.25 |

\*Mean min–max indicates the lowest and highest phenotypic mean among allele-combination classes. Allele combination min–max indicates the allele combination corresponding to the lowest mean followed by the allele combination corresponding to the highest mean. Mean range was calculated as the difference between the highest and lowest allele-combination mean. ProbF values recorded as 0.00E+00 in the original output are displayed as <1.0E–16. The full reciprocal-collapsed epistatic interaction output is provided in Supplementary Table 4, and detailed allele-combination means for the representative interactions are provided in Supplementary Table 5.

For RWC, the leading interaction was detected between TGWA25K-TG0299 on chromosome 3B and TA009869-0749 on chromosome 5D. Across the five representative RWC interactions, allele-combination means ranged from 59.25% to 76.53%, while the within-pair contrasts ranged from 9.07 to 13.76 percentage points. PH showed the largest number of unique marker-by-marker interactions and the strongest phenotypic contrasts among the analyzed traits. The leading representative interaction involved Tdurum_con-tig42008_7461 on chromosome 5D and AX-109475947 in an unassigned genomic region. Across the five representative PH interactions, allele-combination means extended from 88.58 to 130.17 cm, with within-pair contrasts of 34.09–40.40 cm (Table 3). These large differences suggest that two-locus marker combinations may be associated with substantial variation in PH.

Biomass-related traits also showed putative interaction patterns. For DW, the strongest representative interaction was detected between RAC875_c32452_55 on chromosome 1A and RAC875_c16839_188 on chromosome 7B. Across the five representative DW interactions, allele-combination means ranged from 0.10 to 0.19 g, with within-pair contrasts of 0.07–0.08 g. For FW, the leading interaction involved RAC875_c32452_55 on chromosome 1A and AX-158578018 on chromosome 3B. In this trait, allele-combination means ranged from 0.33 to 0.71 g, with within-pair contrasts of 0.28–0.29 g. These results suggest that biomass-related traits may involve non-additive marker effects, particularly because corrected additive SNP associations were not detected for these traits (Table 3).

Flag leaf traits showed more moderate interaction-associated contrasts. The leading FLL interaction was detected between AX-94393315 on chromosome 3A and wsnp_Ex_c1943_3663067 on chromosome 5B. Across the five representative FLL interactions, allele-combination means ranged from 18.59 to 25.80 cm, with within-pair contrasts of 3.62–6.54 cm. For FLW, the strongest interaction involved Tdurum_contig5522_455 on chromosome 5B and BS00049818_51 on chromosome 6D; allele-combination means ranged from 1.60 to 1.86 cm, with contrasts of 0.25–0.26 cm (Table 3). Overall, these results indicate that selected marker-by-marker interactions may contribute to phenotypic variation in water-status, biomass-related, and plant-architecture traits under rainfed conditions.

### 3.7. Candidate gene localization across GWAS-associated genomic regions

Candidate gene localization was performed for GWAS-associated genomic regions identified for the evaluated drought-related traits. Within the ±500 kb physical interval centered on the Bonferroni-significant RWC-associated SNP AX-86184518 (2D: 560,005,079 bp), 12 non-redundant high-confidence genes were identified (Figure 3; Supplementary Table 6). AX-86184518 was located in an intergenic region between TraesCS2D03G1000800 and TraesCS2D03G1001000. The closest gene was TraesCS2D03G1001000, with the nearest gene boundary located only 291 bp from the SNP position. This gene was predicted to encode a 795-aa protein corresponding to UniProt A0A1D5UM13 and RefSeq protein XP_044334216.1. Although annotated as an uncharacterized protein, family-domain evidence indicated similarity to a chloroplastic WEB family-like protein. BLASTp analysis revealed highly conserved cereal homologues, including a Triticum turgidum subsp. durum protein with 97% identity, 98% positives, and an E-value of 0.0. In silico expression profiling of the RefSeq v1.1 counterpart TraesCS2D02G446900 showed detectable expression in both non-stress and abiotic-stress-associated samples, with higher aggregated mean TPM under abiotic conditions than under non-abiotic conditions (12.75 vs. 8.01 TPM; Supplementary Table 7). The neighbouring TraesCS2D03G1000800 gene, located approximately 3.5 kb from AX-86184518, encoded a PEPCK-like protein. Based on physical proximity, conserved WEB family-like features, and detectable expression under abiotic-stress-related conditions, TraesCS2D03G1001000 was retained as the most proximal candidate gene for the RWC-associated 2D locus.

**Figure 3.**
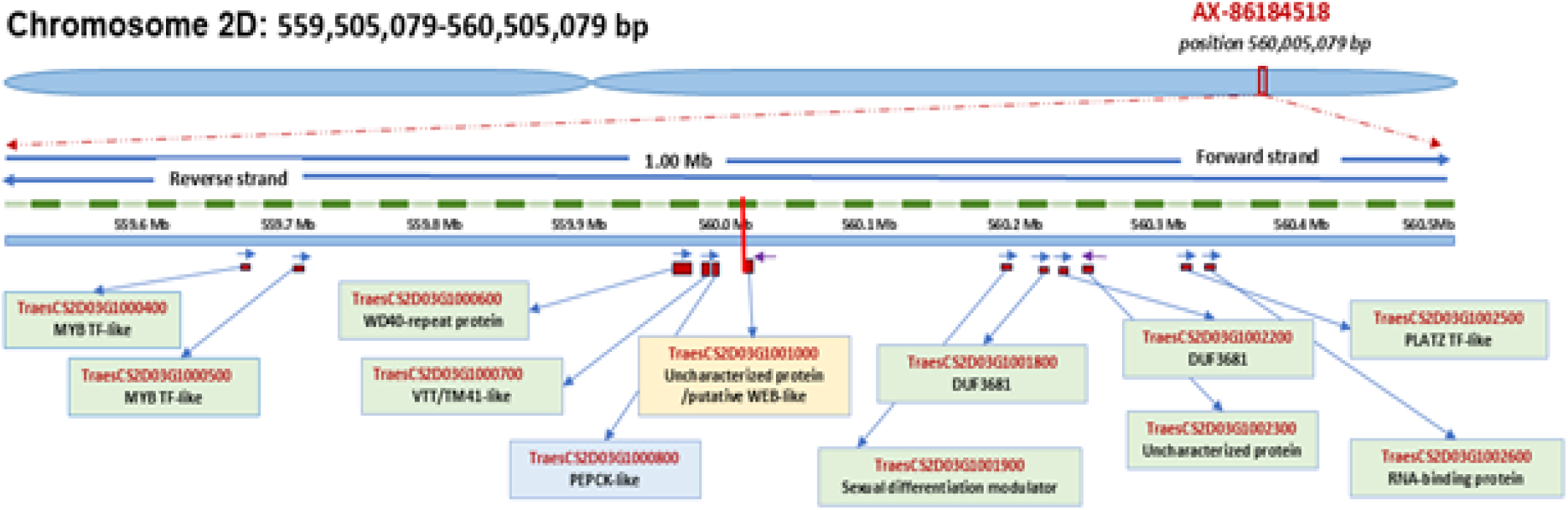
Candidate gene region surrounding the RWC-associated SNP AX-86184518 on chromosome 2D. The schematic shows the ±500 kb interval around the Bonferroni-significant RWC-associated SNP AX-86184518 (2D: 559,505,079–560,505,079 bp) based on the IWGSC RefSeq v2.1 wheat genome annotation. The red vertical line marks the lead SNP position at 560,005,079 bp. Gene models are displayed according to physical position and transcriptional orientation on the forward and reverse strands. TraesCS2D03G1001000, located on the reverse strand, was the closest gene to the marker, whereas TraesCS2D03G1000800 represented the nearest functionally annotated flanking gene.

For PH, candidate gene localization was performed for physically anchored Bonferroni-significant loci. The two closely positioned markers on chromosome 2A, AX-158540611 and AX-158561628, were treated as a single major QTL region containing four non-redundant high-confidence genes. AX-158540611 was located approximately 4.6 kb from TraesCS2A03G1044800, encoding a non-specific lipid-transfer protein/DIR1-like protein, whereas AX-158561628 was positioned close to TraesCS2A03G1044200, annotated as a CobW/HypB/UreG-like nucleotide-binding protein with zinc-regulated GTPase metalloprotein activator domains. The PH-associated region tagged by AX-94514459 on chromosome 4A contained 13 non-redundant genes, including amino acid transporter-like proteins, F-box proteins, lipid-transfer proteins, ribulose-phosphate 3-epimerase, cytochrome P450, pentatricopeptide repeat-containing proteins, and RNA-binding proteins. Additional PH-associated regions on chromosomes 4D, 1D, 1B, and 1A included genes related to transcriptional and post-transcriptional regulation, protein turnover, mitochondrial function, membrane transport, lipid metabolism, and stress-associated protein domains (Supplementary Table 6). The PH-associated SNP on an unassigned contig was excluded because no reliable chromosomal interval could be defined. Candidate genes for selected DW-, FW-, FLL-, and FLW-associated regions were also retrieved and are summarized in Supplementary Table 6.

## 4. Discussion

The present study showed that drought-related traits in the evaluated bread wheat panel were not controlled as a single uniform response, but rather as partly independent physiological and morphological components. This was most clearly evident from the correlation analysis. Biomass traits were strongly associated with flag leaf morphology, whereas RWC showed only a limited association with biomass and almost no relationship with PH. This suggests that the ability to maintain leaf hydration under terminal rainfed stress was not simply a consequence of larger plant size or greater vegetative growth. Similar conclusions have been drawn from physiological studies in wheat, where drought adaptation was shown to depend on coordinated but distinct mechanisms, including water retention, osmotic adjustment, stomatal regulation, and maintenance of photosynthetic activity (Sallam et al., 2019; Guizani et al., 2023).

The partial independence of RWC from plant height and dry biomass is important for interpreting the GWAS results. RWC is commonly considered a direct indicator of leaf water status, but it is not necessarily expected to follow the same genetic architecture as growth-related traits. Condorelli et al. (2022) showed in durum wheat that osmotic adjustment, RWC, and chlorophyll retention were physiologically connected under drought; however, their genetic control was distributed across several genomic regions. Similarly, Czyczyło-Mysza et al. (2018) showed that traits related to excised-leaf water loss are genetically complex and may overlap with leaf morphology without being simply determined by leaf size. The strongest association for RWC was detected on chromosome 2D at AX-86184518. This locus is of particular interest because it passed the corrected significance threshold under rainfed field conditions, where physiological traits are usually affected by environmental heterogeneity. Previous studies have reported loci associated with RWC or water status on other chromosomes, including SSR markers on 5A and 5D in biparental wheat material (Naroui Rad et al., 2012), as well as drought-related loci on 2A identified through association mapping (Ahmad et al., 2014). In contrast, the present association on chromosome 2D appears to represent a different genomic region expressed under terminal field stress. Therefore, AX-86184518 can be considered a promising candidate marker for leaf water-status maintenance in this germplasm. This is relevant because RWC is strongly influenced by developmental stage, stress intensity, and genetic background, as also shown in recent GWAS studies of wheat under drought, osmotic, and salt-related stress (Javid et al., 2022; Reddy et al., 2023; Nouraei et al., 2024). The candidate-gene content around AX-86184518 gives this locus additional biological interest. The nearest gene, TraesCS2D03G1001000, is annotated as an uncharacterized WEB-family-like protein. WEB/PMI2-related proteins have been studied in Arabidopsis as components involved in blue-light-induced chloroplast photorelocation (Luesse et al., 2006; Kodama et al., 2010). This provides a plausible connection with chloroplast-associated stress responses. Under terminal drought, leaves are exposed not only to water deficit, but also to high irradiance and oxidative pressure; therefore, genes related to chloroplast positioning or chloroplast protection represent biologically relevant positional candidates near a water-status locus. The neighbouring gene TraesCS2D03G1000800, annotated as a PEPCK-like protein, is also physiologically relevant. PEPCK-related enzymes participate in organic acid metabolism and carbon remobilization, processes that may influence source–sink balance and stress responses (Lea et al., 2001; Yang et al., 2026). Together, these candidate genes suggest that the 2D region may be connected with chloroplast function, carbon metabolism, and stress-related physiological adjustment, although functional studies will be needed to define their direct roles in wheat drought response.

PH showed a much clearer additive genetic pattern than the other traits. Several markers passed the corrected threshold, especially on chromosomes 2A and 4A, indicating that PH was under strong genetic control in this panel. This is consistent with previous wheat studies, where PH has repeatedly been shown to be a highly heritable trait controlled by loci affecting internode elongation, semi-dwarfing response, and overall plant architecture (Rebetzke and Richards, 2000; Mo et al., 2018; Zhang et al., 2025). In the present study, PH was not significantly correlated with RWC, biomass traits, or flag leaf dimensions. This suggests that, in this germplasm, height variation was largely independent from leaf hydration and vegetative biomass formation. For breeding under rainfed conditions, this is useful because plant architecture may be adjusted without necessarily changing water-status traits. The detected PH loci on 2A and 4A are also interesting in relation to previous mapping studies. Chromosome 2A has repeatedly been implicated in plant-height variation in wheat, including recent analyses of PH-related traits (Zhang et al., 2025). Chromosome 4A has also appeared in studies of drought- and heat-adaptive traits, where QTLs for canopy temperature and yield-related responses under stress were reported (Pinto et al., 2010). Therefore, the present results support the general importance of these chromosomes for wheat architecture and stress-adaptive plant development.

Biomass traits showed a different pattern. Although fresh and dry weight were closely correlated and had relatively high heritability, they did not produce additive associations passing the corrected multiple-testing thresholds. This suggests that biomass accumulation under terminal drought in this panel may not be controlled by one or two large-effect loci. Instead, it may depend on many small-effect loci, environment-sensitive effects, or non-additive interactions. This agrees with the broader GWAS literature on wheat drought response, where terminal drought traits at the reproductive stage often show complex and polygenic inheritance (Reddy et al., 2023; Nouraei et al., 2024). The shared suggestive association on chromosome 7A for FW and DW is therefore worth noting as a putative genomic region contributing to biomass accumulation under rainfed stress.

Flag leaf morphology was strongly associated with biomass at the phenotypic level. This is expected, because the flag leaf is an important photosynthetic source during reproductive development and grain filling. Yang et al. (2016) showed that flag leaf length, width, and area are quantitatively inherited and influenced by water regime, whereas Wang et al. (2022) reported that flag-leaf-related loci may affect yield-related traits. However, in the present study, the GWAS evidence for flag leaf traits remained mainly suggestive, supporting the view that flag leaf architecture is controlled by multiple loci and is sensitive to environmental conditions.

The marker Kukri_c37738_417 on chromosome 1D deserves attention because it appeared as a shared suggestive association for RWC, FW, DW, and FLL. Such a pattern may indicate a genomic region where leaf hydration, leaf expansion, and biomass accumulation are partially connected. Although this marker did not pass the corrected significance thresholds, the 1D region may contain linked or pleiotropic variation affecting several drought-related traits. Further haplotype analysis would help clarify whether this association reflects biological pleiotropy or local linkage disequilibrium.

The epistatic analysis helped to explain why some traits with clear phenotypic variation did not show strong additive GWAS associations. Putative marker-by-marker interactions were detected for all traits, with particularly relevant patterns for RWC and biomass traits. For FW and DW, where corrected additive associations were absent, the epistasis results suggest that allele combinations at two loci may contribute to biomass variation under terminal stress. This agrees with the general view that drought adaptation in wheat includes both additive and non-additive genetic components (Sallam et al., 2019; Gupta et al., 2017). Therefore, the epistatic results complement the additive GWAS findings and provide an additional layer for interpreting the genetic architecture of drought-related traits in this panel.

## 5. Conclusions

Overall, the study reveals three main genetic patterns. First, RWC was associated with a strong candidate locus on chromosome 2D, making AX-86184518 the most promising marker for leaf water-status maintenance in this panel. Second, PH was controlled by several strong additive loci and was largely independent from RWC and biomass traits. Third, biomass and flag leaf traits were phenotypically connected but genetically more complex, with mainly suggestive additive markers and evidence of putative epistatic effects. These results are consistent with the current understanding of drought adaptation in wheat as a multi-component trait, where physiological water status, canopy architecture, and biomass formation should be evaluated together rather than reduced to a single marker or trait. Future studies should focus on testing the stability of the 2D RWC locus and the major PH-associated loci across different years, locations, and drought scenarios. Integrating these loci with grain yield, canopy temperature, osmotic adjustment, root traits, and high-throughput phenotyping indices will help to better understand their contribution to drought adaptation and support their potential use in breeding programs.

**Supplementary Materials:** The online version contains supplementary material available at the link: https://doi.org/10.5281/zenodo.21308148

## Author Contributions

I.H. supervised the project and acquired funding. S.R., I.H. and A.A.N. conceptualized the research idea. S.R., A.J., U.G. and F.Kh. conducted the field trials. J.L., A.A.N. and S.R. performed the statistical analysis. S.R. drafted the original manuscript. I.H., J.L., A.A.N. and SR contributed to the review and editing process. All authors read and approved the final manuscript.

## Data Availability Statement

The data that support the findings of this study are available from Dr. Samira Rustamova upon request.

## Abbreviations

The following abbreviations are used in this manuscript:

ANOVA: Analysis of variance
BLASTP: Basic Local Alignment Search Tool for proteins
Chr: Chromosome
CV: Coefficient of variation
DW: Dry weight
FDR: False discovery rate
FLL: Flag leaf length
FLW: Flag leaf width
FW: Fresh weight
GWAS: Genome-wide association study
H²: Broad-sense heritability
HC: High-confidence
LD: Linkage disequilibrium
MAF: Minor allele frequency
NCBI: National Center for Biotechnology Information
PH: Plant height
PVE: Percentage of genotypic variation explained
QTL: Quantitative trait locus
RWC: Relative water content
SNP: Single-nucleotide polymorphism
TPM: Transcripts per million
TW: Turgid weight

